# *n*-hexane fraction of *Zingiber officinale and Moringa oleifera* interferes with biological parameters of *Callosobruchus chinensis* (L.) (Coleoptera: Bruchidae)

**DOI:** 10.1101/2025.10.07.680921

**Authors:** Oluwasegun John Jegede, Seun Olaitan Oladipupo, Olufunmilayo Eunice Oladipo

## Abstract

The development of insecticide resistance, high cost, misuse, dearth of technical expertise, and restrictive legislation associated with synthetic insecticides have necessitated the development of alternatives. Lessons from plant-insect interactions demonstrate that plant terpenes are worthy probes for insecticidal exploration. Hence, this study screened *n*-hexane fractions of *Zingiber officinale* and *Moringa oleifera* oils as protectant against *Callosobruchus chinensis* and revealed their chemical profiles using Gas Chromatography – Mass Spectrometry (GC-MS). *M. oleifera* (LC_50_; 0.007 µl) was found to be more toxic than *Z. officinale* oil (LC_50_; 0.055 µl) to *C. chinensis*. The oils showed a positive correlation with concentration at 24 h (r = 0.959), 48 h (r = 0.977), 72 h (r = 0.915) and 96 h (r = 0.924). GC-MS revealed 21 and 15 volatile compounds in *Z. officinale* and *M. oleifera* oils, respectively. The most domimant were 5-(1, 5-dimethyl-4-hexenyl)-2-methyl-1,3-Cyclohexadiene (13.64 %) and 8-Octadecenoic acid, methyl ester (34.52 %) in *Z. officinale* and *M. oleifera* oils, respectively. The plant fractions reduced the oviposition potential, egg hatching rate, and adult emergence of *C. chinensis*. Taken together, the results demonstrate possible developmental and inhibitory effects of the oils against *C. chinensis* and points to its possible inclusion in Integrated Pest Management (IPM) practices for *C. chinensis*.

**Highlights:**

➢ *n*-hexane fractions from *Moringa oleifera* seeds are more potent against *Callosobruchus chinensis* than that from *Zingiber officinale* rhizome.
➢ Both oils can disrupt biological parameters and induce mortality of *C. chinensis*.
➢ *M. oleifera* can be useful as grain protectant as it is ubiquitous in sub-saharan Africa, including Nigeria, and has been documented to improve grains’ protein quality in storage.

## 1. Introduction

The geometric increase in human population has made the task of food protection of foremost interest. Globally, post-harvest losses of agricultural produce due to insect infestation is one of the major threats to food protection. In Africa alone, post-harvest losses of legumes (especially peas and beans) due to *Callosobruchus* species have been estimated to be about 45 % (Tapondjou et al., 2002). For example, the pulse beetle, *Callosobruchus chinensis* (L.) (Coleoptera: Bruchidae) begins infestation by laying eggs on cowpeas and pigeon peas in the field and continues its development throughout storage thereby inflicting damage on the grain wholesomeness and marketability (Lale and Kabeh, 2004; Gbaye et al., 2011). Insecticides such as permethrin, phosphine, pirimiphos-methyl, and lindane have been routinely used to protect legumes against *C. chinensis* damage. However, the development of resistance, acceptability, limited availability in certain areas, environmental contamination, and legislative restrictions have necessitated the need for the development of sustainable and eco-friendly alternatives (Adarkwah et al., 2017, Dougoud et al., 2019). At the forefront of these is also a weak and under-resourced capacity to control *C. chinensis* (Chaubley, 2008; Ajayi et al., 2014). Thus, the recent trend in the exploration of plant metabolites (also known as essential oils) as candidates to mitigate against *C. chinensis* attack is unsurprising.

Oil fractions from plants have showed promising bioactivity against insect pests including *C. chinensis* (Ajayi et al., 2017; Oladipo et al., 2019; Ribeiro et al., 2020). Plant oil desirability is due to its non-persistence, biodegradability, and low toxicity (Koul et al., 2008; Oladipupo et al., 2019a). These characteristics explain why essential oils (EOs) are perceived as worthy candidates for inclusion in integrated pest management strategies. Moreover, the ease of registration and ubiquity in tropics, especially to farmers in under-developed countries, make EOs worthy candidates for scientific explorations (Oladipupo et al., 2019b). Earlier studies have shown that the content and concentration of EOs are dictated by plant chemotype and genotype, solvent polarity, and extraction method (Boursier et al., 2011; Gahukar, 2014, Dougoud et al., 2019). Thus, chemical characterization of EOs evaluated as candidates for legume protectants is an important step towards understanding structural-activity relationship and subsequent inclusion/adoption as constituents of commercialized botanicals.

Earlier studies have evaluated the bioactivity of some plant oils against *C. chinensis.* Upadhyay et al. (2006) reported ovipositional deterrence by *Capparis decidua* (Forssk.) against *C. chinensis*. Petroleum ether extract of four vegetable seeds in Cucurbitaceae family were found to be effective protectant against *C. chinensis* (Mishra et al., 2007). In contact and fumigation tests, methanolic fractions from 30 plant species were found effective for managing populations of *C. chinensis* (Isman et al., 2001; Kim et al., 2003). Oladipo et al. (2019) reported that *n*-hexane fractions of *Xylopia aethiopica* (Dunal) and *Senna occidentalis* (L.) were more toxic to *C. chinensis* than acetone fractions. Contrastingly, acetone fractions of oils of both plants displayed higher ovipositional and eclosion deterrence than *n*-hexane fractions (Oladipo et al., 2019). Moreover, *Moringa oleifera* Lam. and *Zingiber officinale* (Roscoe) oils have demonstrated insecticide effects against other insect pests. For example, hydrodistillate of *Z. officinale* are toxic to adult *Tribolium* spp. and larvae of *Plodia interpunctella* (Hubner) (Maedeh et al., 2012; Martynov et al., 2019). Ethanolic fraction of *Z. officinale* demonstrated higher toxicity against *C. chinensis* than *M. oleifera* oil (Ajayi et al., 2017).

To the best of our knowledge, the chemical characterization of *n*-hexane fractions of *Z. officinale* and *M. oleifera,* and bioactivity on aspects of biological parameters of *C. chinensis* have never been evaluated. Therefore, the aims of this study were to: (1) reveal profile of chemical components of *n*-hexane fractions of *Z. officinale* and *M. oleifera* using Gas Chromatography coupled with Mass Spectrometry (GC-MS), (2) evaluate their toxicity against *C. chinensis* over time (hours), and (3) investigate the inhibitory potentials of these oils on oviposition and adult emergence of *C. chinensis*.

## 2. Materials and Methods

### 2.1 Insects

A *C. chinensis* colony was started from an established laboratory colony maintained at the insectaries of the postgraduate research laboratory of the Department of Biology, Federal University of Technology, Akure, Nigeria. The laboratory colony were reared on disinfested cowpea seeds in a 1-liter glass jars at ambient temperature of 28 ± 2 ^°^C, with 75 ± 5 % RH, and a photoperiod of 12:12 (L: D) h. Five pairs of newly emerged adults (0–24 h old) of *C. chinensis* were selected from the laboratory colony and introduced into 250 ml plastic jars containing 200 g of disinfested cowpea seeds. The plastic jars were covered with muslin cloth to allow for ventilation and prevent the escape of *C. chinensis*. Adults (0–24 h old) that subsequently emerged from this were used for the bioassays.

### 2.2 Plant materials and oil extraction

Rhizomes of *Z. officinale* and seeds of *M. oleifera* were purchased from the central market (Oja–Oba) in Akure municipal, Ondo state, Nigeria. Purchased specimens were confirmed by taxonomist and the valid correct names were confirmed at the plant list (available at http://www.theplantlist.org/). The plants were sorted to remove impurities, air dried on a laboratory bench, and ground into a fine powder with a marlex grinder (USHA 500-Watt Motor Power). The powder from each plant were sieved using 16 mesh size sieve. 300 g of each plant powder was soaked in 900 ml *n*-hexane in round-bottomed glass jar for 72 h and periodically stirred. Each solution was sieved using muslin cloth and concentrated using rotary evaporator at 40 ^°^C and rotary speed of 138 to 148 rpm for 3–4 hours. The resulting oil was air-dried to allow for traces of *n*-hexane to escape. The crude oil from each plant was diluted to obtain concentrations of 1-5 % (v/w) with the solvents (*n*-hexane).

### 2.3 Bioassays

#### 2.3.1 Chemical composition analysis of n-hexane fractions of Z. officinale and M. oleifera

Purification of oils were done using liquid-liquid chromatography with sodium sulphate on separating funnel packed with silica gel. The chemical composition analysis of *n*-hexane fractions of *Z. officinale* and *M. oleifera* were carried out using a gas chromatograph (model 7890A; Agilent Technologies, Palo Alto, CA, USA) equipped with an Agilent J & W non-polar HP-5MS capillary column (30 cm X 0.320 mm; 0.25 µm film thickness), mass spectrometer (5975C VLMSD), and an injector (7683B series).

The oven temperature was set at 80 ^°^C for two min, increased by 6 ^°^C per min until a temperature of 240 ^°^C was reached, and held constant at this temperature for 6 min. Helium carries gas flow, which was maintained at 100 kPa. A 1 µl aliquot of each oil was injected in a mass spectrometer. The run time for each sample was 36 min. The peak of each chemical component was expressed based on its retention time and abundance. The identification of components was achieved by searching the mass spectra database and checking for direct similarities with identified components in the system (Adams, 2001). The National Institute of Standards and Technology (NIST) library was also accessed for components’ properties and documentation.

#### 2.3.2 Assessment of the toxicity of n-hexane fractions of Z. officinale and M. oleifera

To evaluate the toxicity of *Z. officinale* and *M. oleifera* oils against *C. chinensis*, five test concentrations (10, 20, 30, 40 and 50 µl v/w) were used. One ml of each oil test concentration was applied to 20 g of cowpea seeds in 250 ml round-bottomed transparent plastic containers. Containers were agitated for 5 - 10 min to ensure uniform coating of the seeds with the oils. Five pairs of 0 – 24 h old *C. chinensis* adults were introduced into each container. Untreated and solvent (*n*-hexane) controls were similarly set up. All experiments were replicated three times in Completely Randomized Design (CRD). Adult mortality was recorded for 96 h at 24 h interval. At the expiration of 96 h; all adult insects, dead or alive, were removed from all replicates.

#### 2.3.3 Ovipositional inhibitory bioassay of n-hexane fractions of Z. officinale and M. oleifera

Number of eggs laid on the seeds from the above set-up in each replicate were recorded. Observation for adult emergence commenced at 26^th^ day post-infestation. Number of adults that emerged from each replicate were recorded. Percent reduction in adult emergence of F_1_ progeny was calculated using:

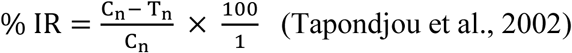

Where C_n_ is the number of emerged insects in the control and T_n_ is the number of emerged insects in the treated containers.

### 2.4 Statistical analysis

Adult mortality data were corrected with mortality obtained in control (untreated) using Abbott formula (Abbott, 1925). Data transformation was done using square-root (for count data) and arc-sine (for percentages) transformation (Tukey, 1977). Transformed data were analyzed using One-way Analysis of Variance (ANOVA). Tukey’s post-hoc test was used to separate the means at α = 0.05. Also, factorial analysis was conducted on mortality data to compare all variable factors for possible interactions using Minitab version 17 (Payton et al., 2006). The median lethal concentration (LC_50_) was calculated using Probit analysis (Finney, 1971). The relationship between concentrations and percentage mortalities, at different exposure periods (24 – 96 h post-treatment) was determined using regression analysis. The corresponding correlation coefficients (r) for prediction equations were also established. All analyses, unless otherwise stated, were done using IBM SPSS software version 20 (IBM SPSS Inc., 2011).

## 3. Results

### 3.1 Acquisition and chemical profile of n-hexane fraction of *Z.* officinale and M. oleifera

The chromatograms showing profiles of the components in *Z. officinale and M. oleifera* oils are shown in **Figures 1** and **2**, respectively. The GC-MS analysis revealed 21 volatile compounds in *Z. officinale* rhizome (**Table 1**). The most dominant were 5-(1, 5-dimethyl-4-hexenyl)-2-methyl-1, 3-Cyclohexadiene (13.64 %) and dec-4-en-3-one-1-(4-Hydroxy-3-methoxyphenyl) (9.57 %). 15 volatile compounds were identified in *M. oleifera* seed (**Table 2**). 8-Octadecenoic acid, methyl ester (34.52 %) and cis-Vaccenic acid (16.11 %) were the dominant components.

**Figure 1.**
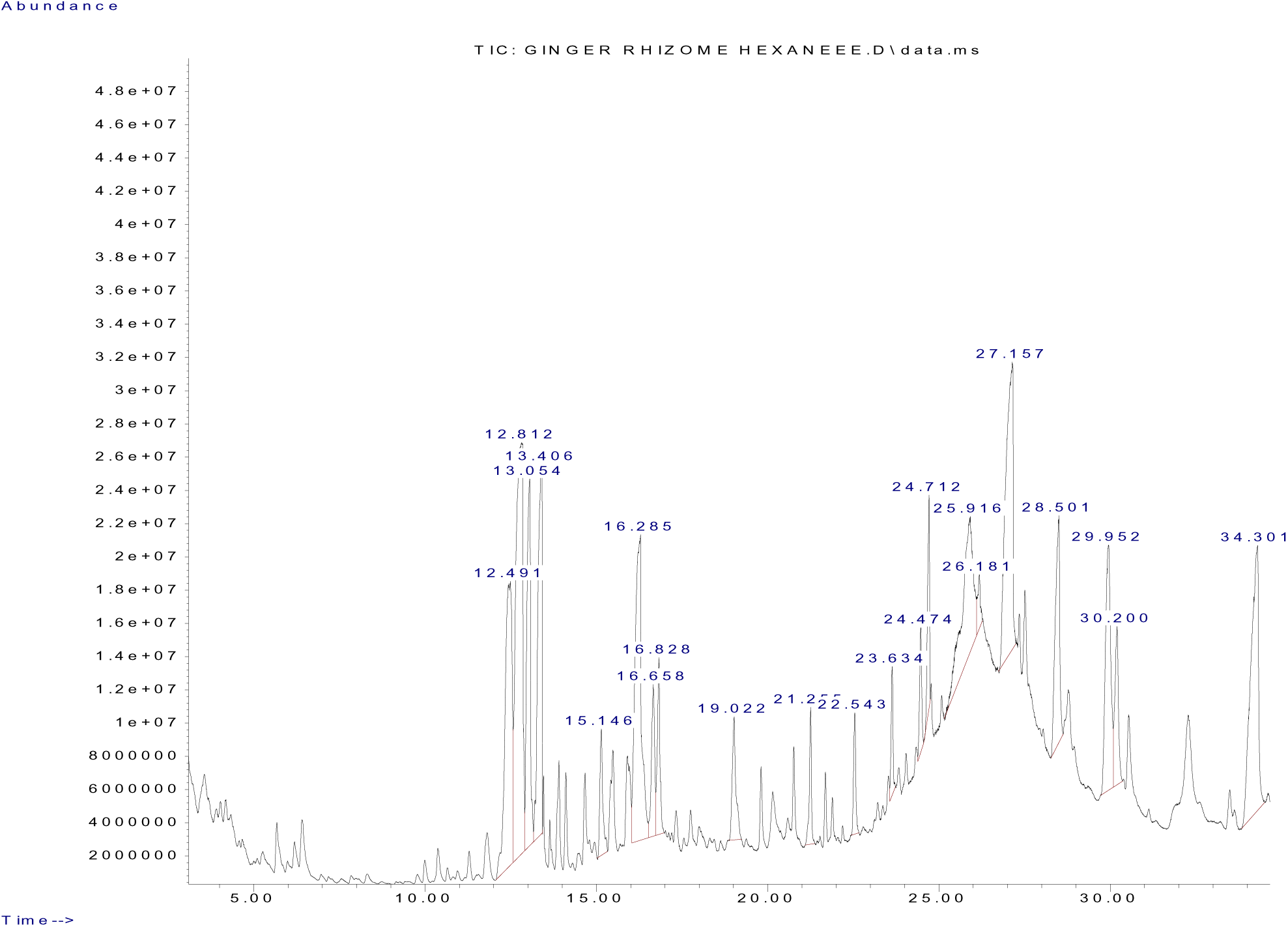
Chromatogram showing the chemical components of *n*-hexane fraction of *Zingiber officinale* rhizome

**Figure 2.**
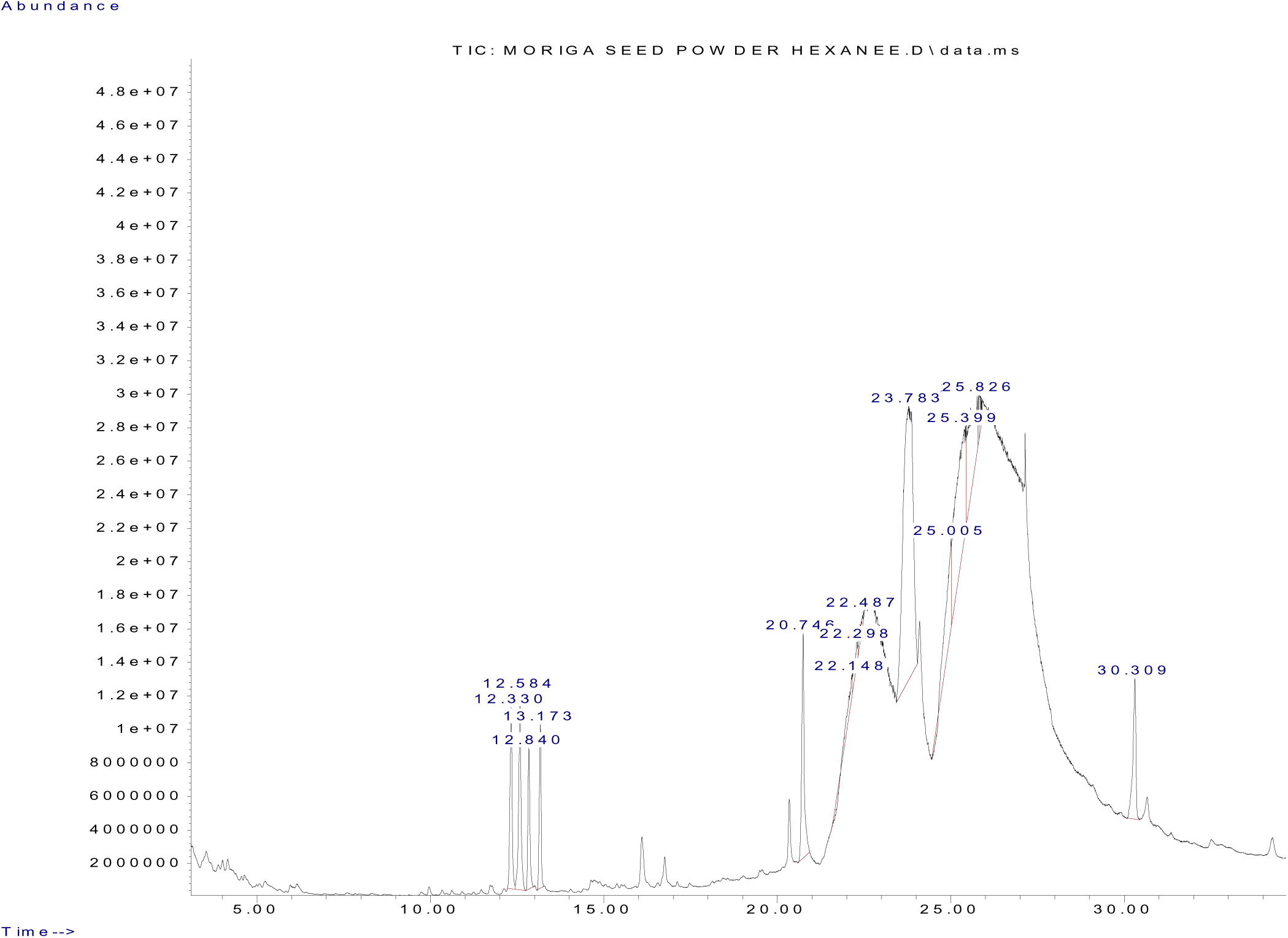
Chromatogram showing the chemical components of *n*-hexane fraction of *Moringa oleifera* seed

**Table 1.** Relative amount and chemical components of *n*-hexane frcation of *Z. officinale rhizome*.

| RT (mins) | Name of components | CAS no. | % of total<br>(Rel. amt.) |
| --- | --- | --- | --- |
| 12.491 | 1-(1, 5-dimethyl-4-hexenyl)-4-methylBenzene | 000644-30-4 | 9.133 |
| 12.812 | 5-(1, 5-dimethyl-4-hexenyl)-2-methyl-1, 3-Cyclohexadiene [S-(R*, S*)] | 000495-60-3 | 13.635 |
| 13.054 | α-Farnesene | 000502-61-4 | 6.365 |
| 13.406 | 3-(1, 5-dimethyl-4-hexenyl)-6-methylene-Cyclohexene [S-(R*, S*)] - | 020307-83-9 | 7.000 |
| 15.146 | (1R, 4R)-1-methyl-4-(6-methylhept-5-en-2-yl)-Cyclohex-2-enol | 058334-55-7 | 1.691 |
| 16.285 | 4-(3-hydroxy-2-methoxyphenyl)-Butan-2-one | 303187-89-5 | 9.241 |
| 16.658 | cis-Z-α-Bisabolene epoxide | 1000131-71-2 | 2.201 |
| 16.828 | 1-Bromo-3, 7-dimethyl-2, 6-octadiene | 035719-26-7 | 2.112 |
| 19.022 | α-2-dimethyl-2-(4-methyl-3-pentenyl)-, [1-α-(R*), 2- $\alpha$]-<br>Cyclopropanemethanol | 121959-70-4 | 2.073 |
| 21.255 | (*E*)-1-(6, 10-dimethylundeca-5, 9-die n-2-yl)-4-methylBenzene | 055968-43-9 | 1.292 |
| 22.543 | 3, 7,11,15-tetramethyl-1,6,10,14-Hexadecatetraen-3-ol (*E,E*) | 001113-21-9 | 1.191 |
| 23.634 | methyl ester, 11-Octadecenoic acid | 052380-33-1 | 1.092 |
| 24.474 | 7-epi-cis-Sesquisabinene hydrate | 1000374-17-9 | 1.200 |
| 24.712 | ethyl ester (*E*)-9-Octadecenoic acid | 006114-18-7 | 1.956 |
| 25.916 | dec-3-en-5-one-(*E*)-1-(4-Hydroxy-3-methoxyphenyl) | 863913-65-9 | 6.909 |
| 26.181 | 2-Vinyl-4, 6-diamino-S-triazine | 003194-70-5 | 0.727 |
| 27.157 | dec-4-en-3-one-1-(4-Hydroxy-3-methoxyphenyl) | 000555-66-8 | 9.574 |
| 28.501 | decan-3-one-5-Hydroxy-1-(4-hydroxy-3-methoxyphenyl) | 039886-76-5 | 4.579 |
| 29.952 | dodec -4-en-3-one-1-(4-Hydroxy-3-methoxyphenyl) | 036700-45-5 | 6.015 |
| 30.200 | decane-3, 5-diyl diacetate-(3R, 5S)-1-(4-Hydroxy-3-methoxyphenyl) | 143615-75-2 | 2.618 |
| 34.301 | tetradec-4-en-3-one-1-(4-Hydroxy-3-methoxyphenyl) | 036752-54-2 | 9.396 |

\*RT – Retention Time
\*CAS no. – Chemical Abstracts Service Registry Number
\* Rel. amt. – Relative amount

**Table 2.**
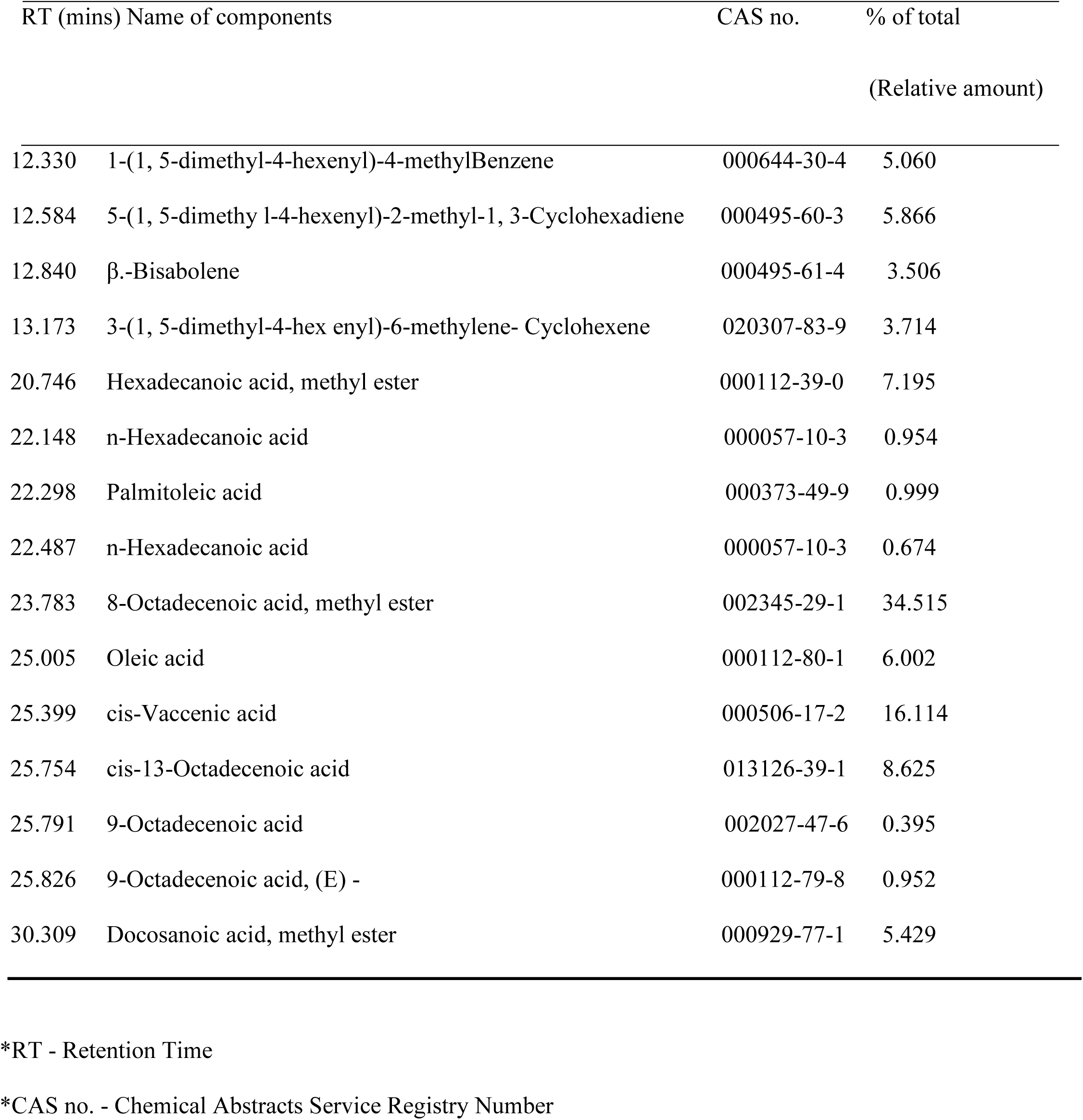
Relative amount and chemical components of *n*-hexane fraction of *M. oleifera* seed.

### 3.2. Toxicity of n-hexane fractions of *Z.* officinale and M. oleifera

There was no mortality in the control experiment at 24 – 72 h for *Z. officinale* oil, and 24 h for *M. oleifera* oil (**Figure 3**). At 24 h, only the highest concentration achieved mortality (3.33 %) of *C. chinensis* for *Z. officinale* oil. The mortality rates of *C. chinensis* ranged from 20 – 40 % at 20 – 50 µl for *M. oleifera* oil. Overall (24 – 96 h), the increase of *M. oleifera* oil concentration led to an increase in the mortality of *C. chinensis*. After 72 h, there was a progression in toxicity of (3.33 – 23.33 %) of *Z. officinale* oil against *C. chinensis*. Statistical analysis revealed no signficant differences (P = 0.141) in mortality between concentrations of *Z. officinale* oil. Mortality rate of *C. chinensis* ranged from 37.04 – 66.67 % for *M. oleifera* oil. The toxicity of *Z. officinale* oil against *C. chinensis* improved after 96 h. However, this was at 50 µl. Relative to *Z. officinale* oil, *M. oleifera* oil exhibited significantly higher toxicity against *C. chinensis* from 24 – 96 h. Thus, median lethal concentration of *M. oleifera* (LC_50_) was much lower (0.007 µl) than that of *Z. officinale* oil (0.055 µl) (**Table 3**). The slope of the log dose-probit line indicated that *C. chinensis* exposed to *M. oleifera* oil had the shallowest slope (0.96) while *C. chinensis* exposed to *Z. officinale* had the steepest slope (2.69) indicating heterogeneous and homogeneous response to both oils, respectively. Regression analysis revealed significant (P < 0.05) positive correlation between mortality and oil concentrations at all exposure periods: 24 h (r = 0.959), 48 h (r = 0.977), 72 h (r = 0.915) and 96 h (r = 0.924) (**Table 4**) of *C. chinensis* exposed to *Z. officinale* and *M. oleifera* oils. Likewise, there were significant impact of the interactions of time X concentration (T X C) (F_10,192_ = 5.95, P < 0.0001).

**Figure 3.**
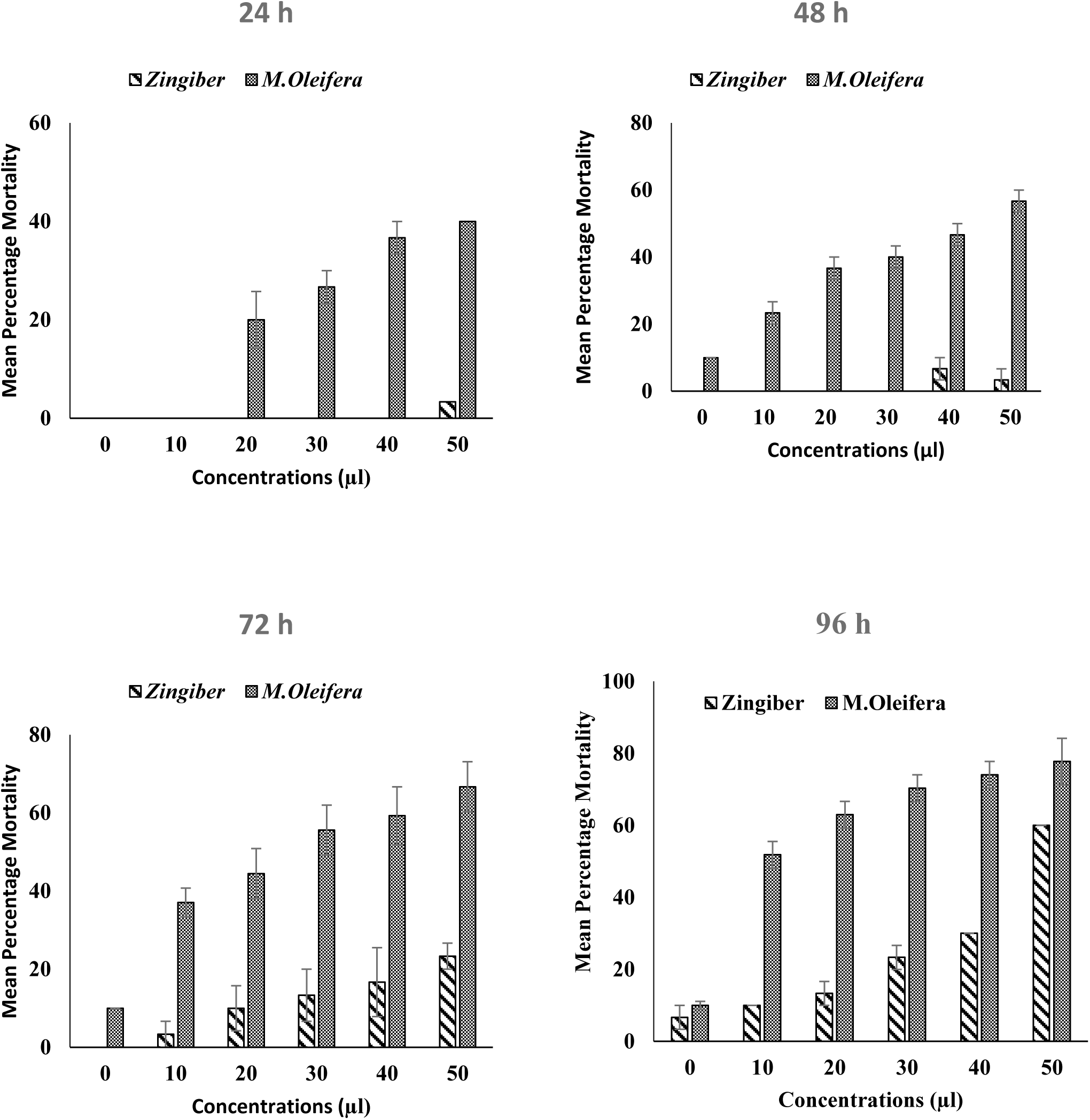
Percentage mortality of *C. chinensis* exposed to *n*-hexane fractions of *Z. officinale* and *M. oleifera*. 0 = control group. Each bar corresponds to the mean ± SE of four replicates.

**Table 3.** Median Lethal Concentration (LC_50_ in ml/20 g of cowpea) of *n*-hexane fractions of *Z. officinale* and *M. oleifera* on *C. chinensis*.

| Plant Material | Slope (±SE) | Intercept (±SE) | $\chi^2$ | df | LC<sub>50</sub> (95 % FL) |
| --- | --- | --- | --- | --- | --- |
| *Z. officinale* | 2.14 (0.54) | 2.69 (0.82) | 5.68 | 1 | 0.055 (0.041 – 0.106) |
| *M. oleifera* | 0.96 (0.44) | 2.08 (0.71) | 2.53 | 1 | 0.007 (0.000 – 0.014) |

$\chi^2$ : Chi-square; SE: Standard error; FL: Fiducial limits; LC: Lethal concentration; df: degree of freedom.

**Table 4.** Relationship between concentration of *n*-hexane fractions of *Z. officinale* and *M. oleifera* and mortality of *C. chinensis*.

| Exposure periods | Correlation coefficient (r) | Regression equation |
| --- | --- | --- |
| **24** | 0.959 | Y = 0.617 + 0.071X |
| **48** | 0.977 | Y = 1.876 + 0.072X |
| **72** | 0.915 | Y = 1.960 + 0.100X |
| **96** | 0.924 | Y = 2.875 + 0.115X |

### 3.3 Oviposition inhibitory assay

There was a significant increase (P < 0.05) in the number of eggs laid by *C. chinensis* on the untreated control and solvent control (**Figure 5**) compared to the plant fractions. The egg counts recorded on seeds treated with *Z. officinale* oil at 20 µl (80.67), 30 µl (80.33) and 40 µl (80.00) are comparable with one another. On the other hand, the egg counts on the seeds treated with *M. oleifera* oil differed significantly (p < 0.05), one from another at the concentrations investigated. Taken together, the lowest number of eggs was observed on cowpea seeds treated with of *M. oleifera* oil at 50 µl (34.67).

**Figure 5.**
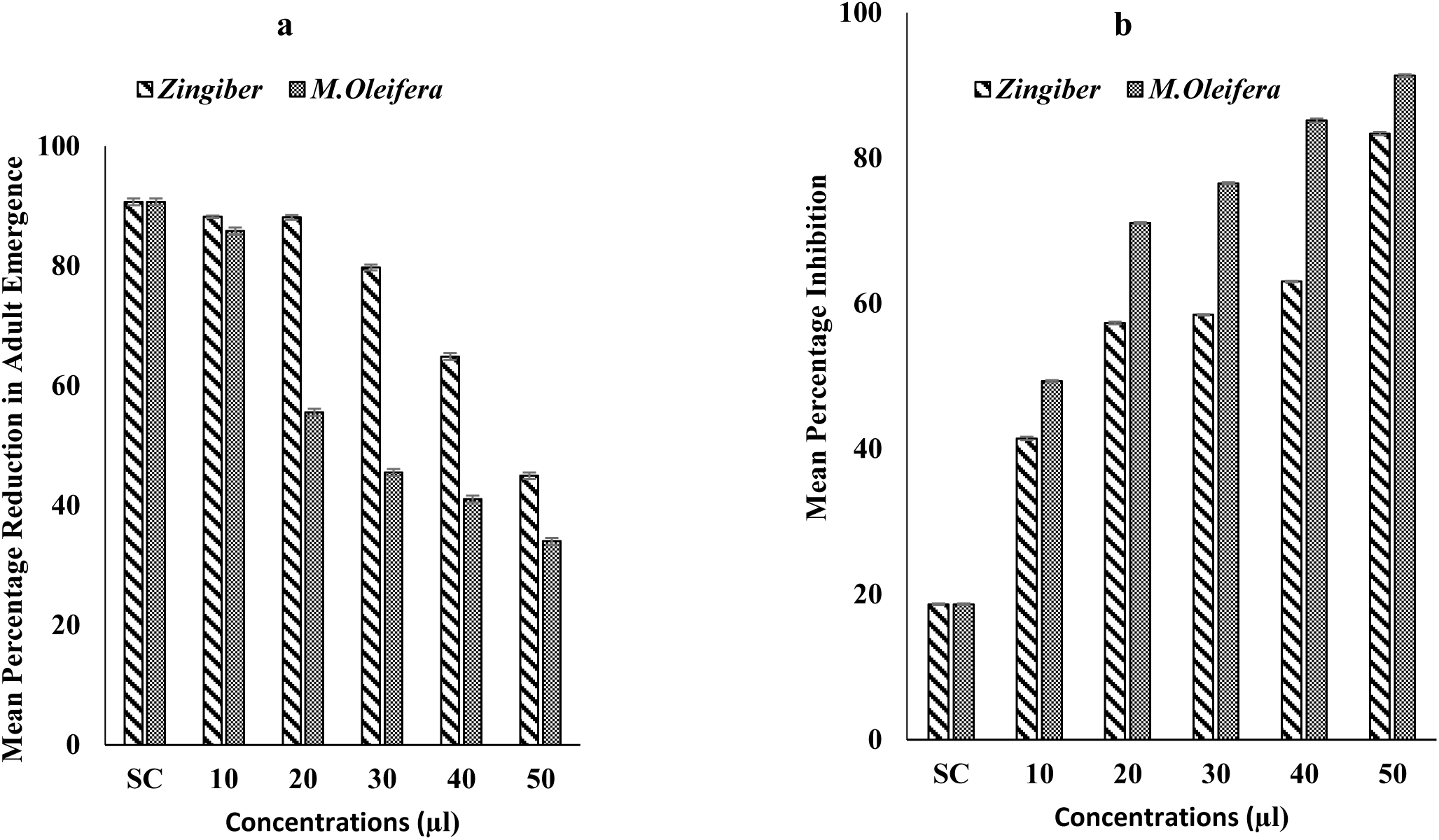
Mean Percentage (a) reduction in adult emergence, and (b) inhibition of *C. chinensis* from eggs laid on cowpeas treated with *n-*hexane fractions of *Z. officinale* and *M. oleifera*. SC = solvent control. Each bar corresponds to the mean ± SE of four replicates.

### 3.3 Variable factors and toxicity of n-hexane fractions of Z. officinale and M. oleifera to C. Chinensis

General Linear Model (GLM) revealed significant differences in the effect of time (T) (F_3,192_ = 191.35, P < 0.0001) and concentration (C) (F_5,192_ = 198.89, P < 0.0001) on the mortality.

Mean percent reduction in *C. chinensis* emergence ranged from 44.95 – 90.71 %, while inhibition ranged from 18.63 – 83.38 % for *Z. officinale* oil. For *M. oleifera* oil, the range of percent reduction in emergence and inhibition were 34.01 – 90.71 % and 18.63 – 91.67 %, respectively.

### 3.4 Developmental inhibitory effects of n-hexane fractions of Z. officinale and M. oleifera

There was similarity in the trend of adult emergence (**Figure 5a**) and mean inhibitory (**Figure 5b**) effects of *Z. officinale* and *M. oleifera* oils. Significantly higher (P < 0.05) number of *C. chinensis* emerged from the control. However, for both oils, a dose-dependent relationship was observed for adults’ emergence and percent inhibition.

## 4. Discussion

The application of insecticides (commonly pyrethroids and organophosphates) for the control of the notorious pest, *C. chinensis*, remains the most popular among farmers and other insect pest control managers. However, due to continued exacerbation of insecticide resistance, high cost, misuse, dearth of technical expertise, and restrictive legislation associated with synthetic insecticides, the continued use cannot be further encouraged. Thus, there is a sought interest in the provision of effective and sustainable insect pest control agents. As alternatives, lessons from plant - insect interactions demonstrate that plant terpenes are worthy probes for insecticidal exploration (Berenbaum et al., 1986; Li et al., 2002). Interestingly, the use of oils from plant materials have shown promising results (Ajayi and Adedire, 2003; Tapondjou et al., 2002, Ajayi et al., 2014; Oladipupo et al., 2019b; Ribeiro et al., 2020). In consonance with this, this research work screened *n*-hexane fractions of *Z. officinale* and *M. oleifera* as protectants against *C. chinensis*. The results demonstrate that the constituent of the plant materials contain chemical compounds with possible inhibitory, neurotoxic, and insecticidal properties.

The present study reports the insecticidal activity of *Z. officinale* and *M. oleifera* oils against *C. chinensis* with *M. oleifera* showing higher toxicity. However, both oils showed a concentration dependent response. Fractions obtained from plant materials have been found to cause severe damage including mortality in *Callosobruchus* species (Mishra *et al*. 2007; Ajayi *et al*. 2014). Chaubley (2008) reported that oils from *Piper nigrum*, *Anethum graveolens*, *Cuminum cyminum and Nigella sativa* caused larvae and adult mortality of *C. chinensis*. In this study, the mortality rate was found to increase with an increase in concentration, and the LC_50_ values decreased at different graded exposure periods indicating that the response of the bruchid to *n*-hexane fractions of *M. oleifera* and *Z. officinale* was concentration and time dependent. The superior toxicity displayed by *M. oleifera* over *Z. officinale* is consistent with the findings of Ajayi et al. (2007).

Moreover, earlier attempt to describe the toxicity of *M. oleifera* against insect pests appear widespread in literature. For example, the presence of the fatty acids 8-Octadecenoic acid, methyl ester and cis-Vaccenic acid in *M. oleifera* has been implicated to cause mortality (Ashfaq *et al*. 2012; Leone *et al*. 2015). These fatty acid cause severe asphyxiation, thereby making breathing impossible in insect (Olayemi and Alabi 1994; Ajayi et al., 2018). Compared to *Z. officinale,* the proportion of fatty acids in *M. oleifera* is about 30 % more. So, this may demonstrate a scientific rationale for the superior toxicity of *M. oleifera*. *Z. officinale* showed delayed toxicity in this study. This contrasts with the findings of Chaubey (2013). Since the bioactivity of a plant metabolite often varies with respect to edaphic and environmental conditions (Gahukar, 2014, Dougoud et al., 2019), the absence of the phytochemical constituent analysis in the study complicates straight forward comparison.

In general, essential oils are sought after because of their biodegradability (high volatility as a funcion of low-persistence in the enviroment). Volatility is a function of hydrogenated to oxygenated ratio with hydrogenated compounds being more volatile (Ahn et al., 2008; Regnault-Roger et al., 2008). Interestingly, the GC-MS results showed that both *Z. officinale* and *M. oleifera* oils have a higher proportion of hydrogenated compunds. These hydrogenated compounds (broadly termed monoterpenoids) have been described to disrupt metabolic pathways, alter biological parameters, and cause death in insects, including *C. chinensis* (Kéita et al., 2001; Jiang et al., 2012; Ribeiro et al., 2020). Thus, the toxicity associated with the oils may be related to their components. Nonetheless, a straight forward assumption that the dominant compounds may have been solely responsible for the observed toxicity should be made with caution as it has been shown that that is not often the case (Veras et al., 2012; Tak and Isman, 2015).

However, essential oils do not only affect insects by causing mortality, but also interferes with life-history traits and biochemical functions. For example, the oils inhibited ovipositional ability of *C. chinensis*, and retarded subsequent morph into adults. Similar observations have been made on eggs of the grain moth and *C. chinensis* (Ileke, 2013; Chaubey, 2013). Interestingly, recent findings have shown that essential oils, including *M. oleifera*, not only represent safe protectants against insect attack, but these oils can also boost the nutritional quality by increasing the amino acid index, protein efficiency ratio, biological value and net protein value (Ilesanmi and Gungula, 2016). More so, synthetic insecticides such as organophosphates routinely used as grain protectants depletes the nutritional constituents of grains in storage, rendering them unsuitable for consumption (Akami et al., 2017).

In conclusion, the findings of this study demonstrate that *n*-hexane fraction of *M. oleifera* are more toxic to *C. chinensis* than *Z. offcinale*. More so, both plant oils can reduce egg hatchability, retard *C. chinensis* development and induced mortality. Taken together, the results demonstrate possible developmental and inhibitory effects of the oils against *C. chinensis* and points to its possible inclusion in Integrated Pest Management (IPM) practices for *C. chinensis.* The ubiquity of these plant material to local farmers can fast-track adoptions in localities where there is widespread insecticide development, restrictive legislation, and lack of funds. However, further study focusing on the mode of action of these oils is merited, before subsequent incorporation into management system for grain (cowpea) protection.

## Acknowledgments

The authors are grateful to Dr. Rotimi Aladesanwa for assistance with data analysis, and Mrs. Toyin Ojo for technical support.

## Author contributions

**Oluwasegun J. Jegede**: Conceptualization, Methodology, Investigation, Data curation, Data analysis, Resources, Writing – review & editing. **Seun O. Oladipupo:** Data curation, Data analysis, Writing – original draft, review & editing. **Olufunmilayo E. Oladipo:** Conceptualization, Methodology, Validation, Supervision, Writing – review & editing.

## References

Abbott, W.S. 1925. A method for computing the effectiveness of an insecticide. J. Econ. Entomol. 18, 265–267.

Adams, R.P. 2001. Identification of essential oil components by gas chromatography – mass spectroscopy. Allured Publishing Co. Carol Stream, IL. 804 p.

Adarkwah, C., Obeng-Ofori, D., Hörmann, V., Ulrichs, C., Schöller, M. 2017. Bioefficacy of enhanced diatomaceous earth and botanical powders on the mortality and progeny production of *Acanthoscelides obtectus* (Coleoptera: Chrysomelidae), *Sitophilus granarius* (Coleoptera: Dryophthoridae) and *Tribolium castaneum* (Coleoptera: Tenebrionidae) in stored grain cereals. Int. J. Trop. Insect Sc. 37, 243–258.

Ajayi, O.E., Appel, A.G., Fadamiro, H.Y. 2014. Fumigation toxicity of essential oil monoterpenes to *Callosobruchus maculatus* (Coleoptera: Chrysomelidae: Bruchinae). J. Insect. Sci Article ID 917212.

Ajayi, O.E., Oladipupo, S.O., Jegede, O.J., 2018. Comparative and synergistic influence of extracts of two tropical plants on the activity of the cowpea weevil, Callosobruchus chinensis. Med. Plant. Res. 8, 60–63.

Akami, M., Chakira, H., Andongma A.A., Khaeso, K., Gbaye, O.A., Nicolas, N.Y., Nukenine, E.N., Niu, C.Y. 2017. Essential oil optimizes the susceptibility of C*allosobruchus maculatus* and enhances the nutritional qualities of stored cowpea *Vigna unguiculata* R. Soc. Open. Sci. 4, 170692.

Ashfaq, M., Basra, S.M., Ashfaq, U. 2012. *Moringa*: A Miracle Plant for Agro-forestry. J. Agr. Forest. Soc. Sci, 8, 115–122.

Berenbaum, M.R., Zanger, A.R., Nitao, J.K. 1986. Constraints on chemical coevolution: wild parsnips and the parsnip webworm. Evol. 40: 1215.

Boursier, C., Bosco, D., Coulibaly, A., Negre, M., 2011. Are traditional neem extract preparations as efficient as a commercial formulation of azadirachtin? Crop Protection 30, 318–322.

Chaubley, M.K. 2008. Fumigant toxicity of essential oils from some common spices against pulse beetle, *Callosobruchus chinensis* (Coleoptera: Bruchidae). J. Oleo Sci. 57, 171–179.

Chaubey, M.K. 2013. Biological activities of *Zingiber officinale* (Zingiberaceae) and *Piper cubeba* (Piperaceae) essential oils against pulse beetle, *Callosobruchus chinensis* (Coleoptera: Bruchidae). Pakistan. J. Biol. Sci.16, 517–523.

Dougoud, J., Toepfer, S., Bateman, M., Jenner, W.H. 2019. Efficacy of homemade botanical insecticides based on traditional knowledge. A review. Agron. Sustain. Dev. 37, 5–22.

Finney, D.J. 1971. Probit Analysis. Cambridge University Press, Cambridge, London, 333 p.

Gahukar, R. 2014. Factors affecting content and bioefficacy of neem (*Azadirachta indica* A. Juss.) phytochemicals used in agricultural pest control: A review. Crop. Prot. 62, 93–99.

Gbaye, O.A., Millard, J.C., Holloway, G.J. 2011. Legume type and temperature effects on the toxicity of insecticide to the genus *Callosobruchus* (Coleoptera: Bruchidae). J. Stored. Prod. Res. 47, 8–12.

IBM SPSS Inc. 2011. Statistical Package for Social Sciences. SPSS, Inc., Chicago, IL.

Ileke, K.D. 2013. Insecticidal activity of four medicinal plant powders and extracts against Angoumois grain moth, *Sitotroga cerealella* (Olivier) [Lepidoptera: Gelechidae]. Egypt. J. Biol. 15, 7–21.

Ilesanmi, J., Gungula, D. 2016. Amino acid composition of cowpea grains preserved with mixtures of neem (*Azadirachta indica*) and moringa (*Moringa oleifera*) seed oils. Am. J. Food. Nutr. 4, 150–156.

Jiang, C.H., Liu, Q.Z., Du, S.S., Deng, Z.W., Liu, Z.L. 2012. Essential oil composition and insecticidal activity of *Evodia lepta* (Spreng)Merr. root barks from China against two grain storage insects. J. Med. Plant. Res. 6, 3464–3469.

Kéita, S.M., Vincent, C., Schmit, J.P., Arnason, J.T., Bélanger, A., 2001. Efficacy of essential oil of *Ocimum basilicum* L. and *O. gratissimum* L. applied as an insecticidal fumigant and powder to control *Callosobruchus maculatus* (Fab.) [Coleoptera: Bruchidae]. J. Stored. Prod. Res. 37, 339–349.

Kim, S.I., Roh, J.Y., Kim, D.H., Lee, H.S., Ahn, Y.J. 2003. Insecticidal activities of aromatic plant extracts and essential oils against *Sitophilus oryzae* and *Callosobruchus chinensis*. J. Stored. Prod. Res. 39, 293–303.

Koul, O., Walia, S., Dhaliwal, G.S. 2008. Essential oils as green pesticides: potential and constraints. Biopestic. Int. 4, 63–84.

Lale N.E.S., Kabeh, J.D. 2004. Pre-harvest spray of neem (*Azadirachta indica* A. Juss) seed products and pirimiphos-methyl as a method of reducing field infestation of cowpeas by storage bruchids in the Nigerian sudan savanna. Int. J. Agric. Biol. 6, 987–993.

Leone, A., Alberto, S., Alberto, B., Alberto, S., Junior, A., Simona, B. 2015. An overview: cultivation, genetic, ethnopharmacology, phytochemistry and pharmacology of *Moringa oleifera* leaves. Int. J. Mol. Sci. 16, 12791–12835,

Li, L., Li, C., Lee, G.I., Howe, G.A. 2002. Distinct roles for jasmonate synthesis and action in the systemic wound response of tomato. Proc. Natl. Acad. Sci. 99, 6416–6421.

Maedeh, M., Hamzeh, I., Hossein, D., Majid, A., Reza, R.K. 2012. Bioactivity of essential oil from *Zingiber officinale* (Zingiberaceae) against three stored-product insect species. J. Essent. Oil. Bear. Pl. 15, 122–133.

Martynov, V.O., Titov, O.G., Kolombar, T.M., Brygadyrenko, V.V. 2019. Influence of essential oils of plants on the migration activity of *Tribolium confusum* (Coleoptera, Tenebrionidae). Biosyst. Divers. 27, 177–185.

Mishra, D., Shukla, A.K., Singh, A., Dixit, A.K., Singh, K. 2007. Efficacy of application of vegetable seed oils as grain protectant against infestation by *Callosobruchus chinensis* and its effect on milling fractions and apparent degree of dehusking of legume-pulses. J. Oleo. Sci. 56, 1–7.

Oladipo, O.E., Oyeniyi, E.A., Aribisala, T.E. 2019. Toxicity and biochemical mechanisms underlying the insecticidal efficacy of two plant extracts *on Callosobruchus chinensis* (Coleoptera: Chrysomelidae) infesting cowpea seeds. J. Crop. Prot. 8, 259–274.

Oladipupo, S.O., Callaghan, A., Holloway, G.J., Gbaye, O.A. 2019a. Variation in the susceptibility of *Anopheles gambiae* to botanicals across a metropolitan region of Nigeria. PLoS One. 14, e0210440.

Oladipupo S.O., Hu, X.P. Appel, A.G. 2019b. Topical toxicity profiles of some aliphatic and aromatic essential oil components against insecticide-susceptible and resistant strains of German cockroach (Blattodea: Ectobiidae), J. Econ. Entomol. xx, 1–9.

Olayemi, A.B., Alabi, R.O. 1994. Studies on traditional water purification using *Moringa oleifera* Seeds. Afr. Study. Monogr. 15, 135–142.

Payton, M.E., Richter, S.J., Giles, K.L., Royer, T.A. 2006. On transformations of count data for tests of interaction in factorial and split-plot experiments. J. Econ. Entomol. 99, 1002–1006.

Regnault-Roger, C., Philogène, B.J.R. 2008. Past and current prospects for the use of botanicals and plant allelochemicals in integrated pest management. Pharm. Biol, 46, 41–52.

Ribeiro, I.A.T.D, Silva, R.D., Silva, A.G.D., Milet-Pinheiro, P., Paiva, P.M.G., Daniela Maria Do Amaral Ferraz Navarro, Silva, M.V.D., Napoleão, T.H., Correia, M.T.D.S. 2020. Chemical characterization and insecticidal effect against *Sitophilus zeamais* (maize weevil) of essential oil from *Croton rudolphianus* leaves. Crop Prot. 29, 105043.

Tak, J.H., Isman, M.B., 2015. Enhanced cuticular penetration as the mechanism for synergy of insecticidal constituents of rosemary essential oil in *Trichoplusia ni*. Sci. Rep. 5, 12690.

Tapondjou, L.A., Alder, A., Bonda, H., Fontem, D.A. 2002. Efficacy of powder and essential oil from *Chenpodium ambrosioides* leaves as post-harvest grain protectants against six stored products beetles. J. Stored. Prod. Res. 38, 395–402.

Tukey, J.W. 1977. Exploratory data analysis. Addison-Wesley, Reading, MA.

Upadhyay, R.K., Mukesh, L.R., Chaubey, K., Subhash, C. 2006. Ovipositional responses of the pulse beetle, *Bruchus chinensis* (Coleoptera: Bruchidae) to extracts and compounds of *Capparis deciduas*. J. Agric. Food Chem. 54, 9747–9751.

Veras, H.N., Rodrigues, F.F., Colares, A.V., Menezes, I.R., Coutinho, H.D., Botelho, M.A., Costa, J.G. 2012. Synergistic antibiotic activity of volatile compounds from the essential oil of *Lippia sidoides* and thymol. Fitoterapia 83, 508–512.

